# In silico discovery of potential inhibitors targeting the MEIG1-PACRG complex for male contraceptive development

**DOI:** 10.1101/2024.12.16.628759

**Authors:** Timothy Hasse, Zhibing Zhang, Yu-ming M. Huang

## Abstract

The interaction between meiosis-expressed gene 1 (MEIG1) and Parkin co-regulated gene (PACRG) is a critical determinant of spermiogenesis, the process by which round spermatids mature into functional spermatozoa. Disruption of the MEIG1-PACRG complex can impair sperm development, highlighting its potential as a therapeutic target for addressing male infertility or for the development of non-hormonal contraceptive methods. This study used virtual screening, molecular docking, and molecular dynamics (MD) simulations to identify small molecule inhibitors targeting the MEIG1-PACRG interface. MD simulations provided representative protein conformations, which were used to virtually screen a library of over 800,000 compounds, resulting in 48 high-ranking candidates for each protein. PACRG emerged as a favorable target due to its flexible binding pockets and better docking scores compared to MEIG1. Key binding residues with compounds included W50, Y68, N70, and E74 on MEIG1, and K93, W96, E101, and H137 on PACRG. MD simulations revealed that compound stability in MEIG1 complexes is primarily maintained by hydrogen bonding with E74 and π-π stacking interactions with W50 and Y68. In PACRG complexes, compound stabilization is facilitated by hydrogen bonding with E101 and π-π interactions involving W96 and H137. These findings highlight distinct molecular determinants of ligand binding for each protein. Our work provides mechanistic insights and identifies promising compounds for further experimental validation, establishing a foundation for developing MEIG1-PACRG interaction inhibitors as male contraceptives.

## Introduction

Spermiogenesis, the final stage of sperm development, involves the transformation of round spermatids into mature spermatozoa. This complex process includes critical changes, such as nuclear elongation and condensation, acrosome formation, and flagellum development, all of which are essential for male fertility^1,2^. A temporary structure known as the manchette plays a pivotal role in shaping the sperm head and transporting proteins required for tail formation^3^. Two proteins, meiosis-expressed gene 1 (MEIG1) and Parkin co-regulated gene (PACRG), have been identified as crucial components of this process^4^. MEIG1 binds to PACRG to form a complex vital for manchette function, and disruption of this interaction leads to defective spermiogenesis and male infertility^5,6^. These findings position the MEIG1-PACRG complex as a promising target for male contraceptive development^7^.

The MEIG1 protein, first discovered in a gene screen related to meiosis, is highly active in the testis and plays an essential role in spermiogenesis^8^. MEIG1 has a unique structural fold composed of alternating α-helices and β-sheets and interacts with PACRG through a well-defined interface involving key residues such as W50 and Y68^6,9^ (Figure 1). PACRG stabilizes microtubules, serving a critical function in organizing the manchette and facilitating sperm tail development^10,11^. Experimental studies have demonstrated that mutations in MEIG1, such as W50A and Y68A, significantly weaken its binding to PACRG, disrupting the manchette’s function and impairing sperm formation^6^.

**Figure 1.**
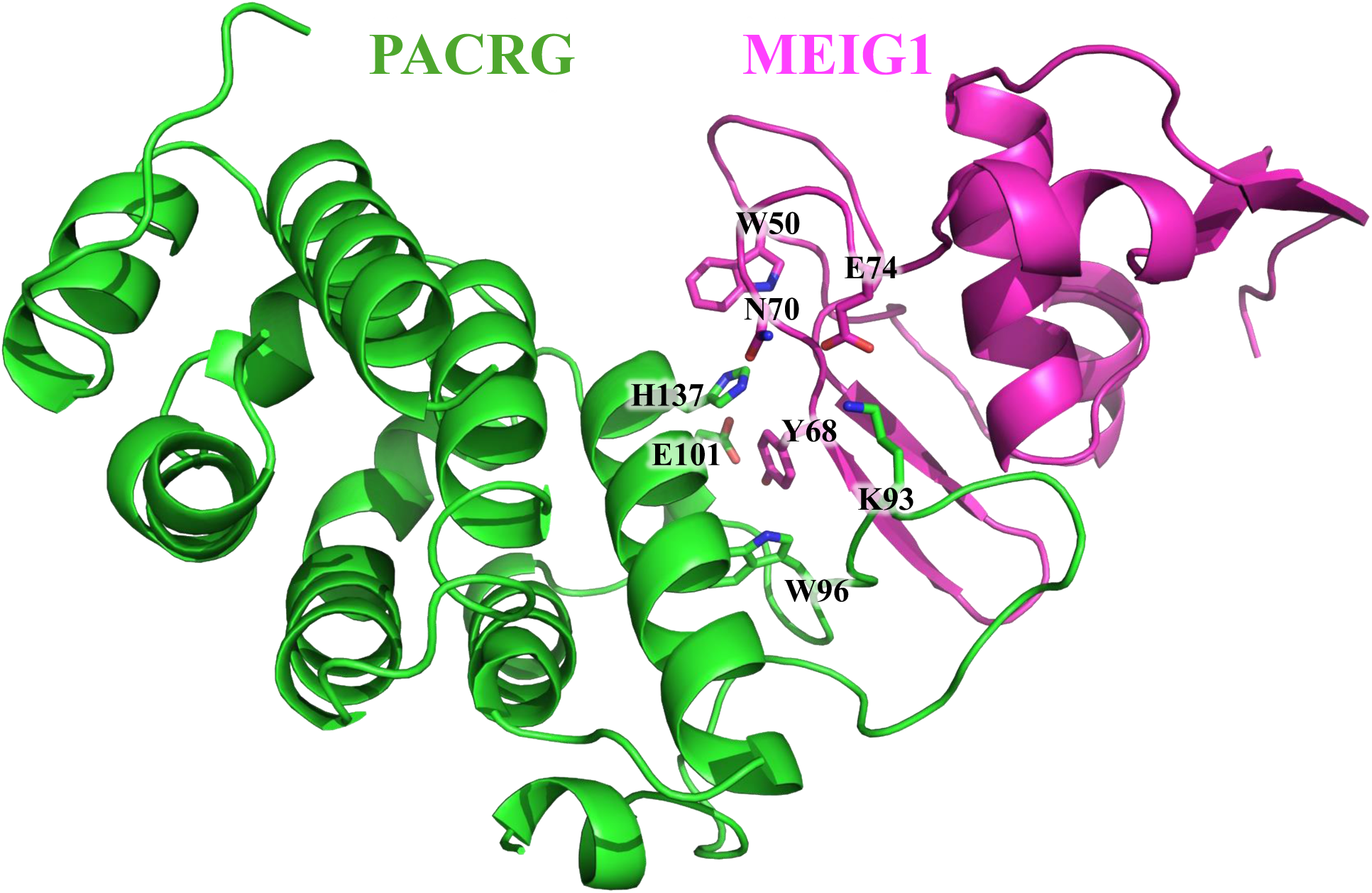
Structure of MEIG1-PACRG complex with MEIG1 in magenta and PACRG in green. Key residues W50, Y68, N70, and E74 of MEIG1 and K93, W96, E101, and H137 of PACRG are displayed.

A recent computational study^12^ explored the dynamic properties of the MEIG1 and PACRG proteins using molecular dynamics (MD) simulations. This research revealed that the W50A and Y68A mutations destabilize the MEIG1-PACRG interface by altering the stability of key residues and disrupting hydrogen bonding networks, including the Y68-E101 hydrogen bond. The study also identified potential ligand binding pockets on both proteins at their interaction interface. PACRG, in particular, emerged as a promising drug target due to the stability and accessibility of its binding pockets.

Building on these findings, this study aims to identify small molecule inhibitors of the MEIG1-PACRG interaction using *in silico* approaches, including virtual screening, molecular docking, and MD simulations. Here, we use representative protein conformations derived from MD simulations to perform virtual screening on over 800,000 compounds for their potential to bind to the MEIG1 and PACRG proteins. By evaluating the binding affinities and stability of these compounds through MD simulations, we aim to identify top candidates capable of disrupting the MEIG1-PACRG interaction and elucidate their binding mechanisms. These results will lay the groundwork for experimental validation and provide valuable insights into the molecular basis of this interaction, advancing the development of a novel male contraceptive.

## Materials and methods

### Obtaining representative conformations of MEIG1 and PACRG

We obtained the MD trajectories of human apo-MEIG1, apo-PACRG, and the MEIG1-PACRG complex from a previous computational study^12^. To identify representative structures from these simulations, we performed clustering based on the root mean square deviation (RMSD) of each frame. RMSD was calculated on the backbone atoms of each frame relative to the initial structure, generating an RMSD matrix. Initially, we preprocessed the trajectories by removing water molecules and ions to focus solely on the protein structures. For the MEIG1-PACRG complex simulations, we generated two separate trajectories from each simulation: one for holo-MEIG1, containing only the MEIG1 protein, and another for holo-PACRG, containing only the PACRG protein. This allowed us to analyze the two proteins individually.

We applied the GROMOS gmx cluster module^13,14^ to cluster the MD trajectories. The cutoff values were adjusted between 1.4 Å to 2.9 Å to obtain five distinct representative protein conformation clusters. For each cluster, we selected the frame with the median RMSD as the representative structure to ensure these structures accurately reflect the simulation trajectory. Additionally, we included the original equilibrated crystal structure as a sixth model for each protein to serve as a reference. These representative structures were subsequently used for virtual screening and molecular docking.

### Virtual screening and molecular docking

We performed virtual screening and molecular docking using the Schrödinger Suite 2021^15–17^ to evaluate the binding affinity of various ligands to the selected binding sites on the MEIG1 and PACRG proteins. The ChemBridge CORE Library^18^, consisting of nearly 800,000 compounds, was chosen for this screening due to its extensive and diverse range of potential small molecule inhibitors.

To prepare the compound library, we used the LigPrep module in Schrödinger^19^ to convert the compounds from 2D to 3D formats, generate multiple poses, add missing hydrogen atoms, adopt correct ionization states, and minimize the structures. This step increases the number of possible poses for each compound to enhance the accuracy of the screening process. We prepared the protein structures of MEIG1 and PACRG from clustering using the Protein Preparation Wizard^20,21^. This involved assigning correct protonation states for biological pH, assigning bond orders, and minimizing the structures using the OPLS_2005 force field^22^.

Receptor Grid Generation^15–17,23^ was then performed to define the docking space around the primary binding pockets identified by the previous computational study. These pockets were centered on key residues involved in the MEIG1-PACRG interface, specifically W50 and Y68 on MEIG1, and E101 and H137 on PACRG. The grid inner box size was set to 10x10x10 Å, and the outer box to 30x30x30 Å, to ensure precise docking within the targeted pocket.

The virtual screening process involved three stages: high throughput virtual screening (HTVS), standard precision (SP), and extra precision (XP)^15–17^. Each stage increased in accuracy and computational demand, progressively filtering the compounds. The cutoff values for HTVS, SP, and XP were set to 0.5%, 2%, and 25%, respectively, which effectively reduced the initial 800,000 compounds to the top candidates. Compounds were ranked based on their docking scores. Visual Molecular Dynamics (VMD)^24^ was used to visualize the protein-ligand complexes and analyze detailed interactions between the top candidates and the proteins.

### Molecular dynamics simulations of key protein-compound complexes

To assess the dynamics of key protein-compound complexes identified from virtual screening and molecular docking, we performed conventional MD simulations on the 10 MEIG1-compound and PACRG-compound complexes that had the best compound binding scores. All MD-based simulations were performed using the AMBER20 package^25,26^. The AMBER ff14SB force field^27^ was used for the protein, and the GAFF2 force field^28^ was used for the compounds. The protein models were minimized in three steps: 500 steps of hydrogen minimization, 5,000 steps of hydrogen and sidechain minimization, and 5,000 steps of whole protein minimization. Next, the complexes were solvated with the TIP3P water model^29^, extending 12 Å from the protein surface, and Cl^−^ were added to neutralize the system’s charge. The system underwent further minimization with 1,000 steps on the water molecules and 5,000 steps on the entire system. A 9 Å cutoff was applied during minimization, and the particle mesh Ewald summation^30^ was utilized for long-range electrostatics.

Following minimization, the systems were gradually heated from 50 K to 300 K in increments of 50 K. Each heating stage lasted 10 ps, except for the final stage at 300 K, which lasted 100 ps. After equilibration, a short 20 ns conventional MD simulation was performed at 300 K to ensure that the systems converged to the appropriate thermodynamic state. All simulations employed Langevin dynamics^31^ with a collision frequency of 5 ps^-1^ in an isothermal-isobaric (NPT) ensemble. Bonds involving hydrogen atoms were constrained using the SHAKE algorithm^32^, and the simulation time step was set to 2 fs. Following this initial MD simulation, a 100 ns conventional MD production run was performed for each complex to analyze the stability of the protein-ligand interactions. The trajectories were collected every 10 ps for analysis.

### Post-molecular dynamics analysis

Visualization of MD trajectories and atom-atom distance measurements for specific interactions were conducted using VMD^24^. The CPPTRAJ program^33^ from the AMBER package^25^ was used to calculate RMSD, hydrogen bond occupancy, and aromatic centroid-centroid distances and planar angles. RMSD was calculated for compound heavy atoms relative to the equilibrated initial docking pose. Hydrogen bonds were defined by a donor-acceptor distance of ≤ 3.5 Å and a donor-hydrogen-acceptor angle of ≥ 120°. Centroid-centroid distances and planar angles were calculated to quantify aromatic interactions between compound aromatic rings and key protein residues. Centroid-centroid distances were measured between the geometric centers of the aromatic rings, and interplanar angles were calculated using the normal vectors of the ring planes. Angles were normalized to 0°–90°. Face-to-face stacking was characterized by centroid-centroid distances of ∼4 Å and planar angles between 0° and 30°, while edge-to-face stacking showed centroid-centroid distances of ∼6 Å and planar angles between 60° and 90°^34–36^.

## Results and discussion

### In silico virtual screening to identify potential MEIG1 and PACRG inhibitors

To identify potential inhibitors of the MEIG1-PACRG complex, we first obtained representative protein conformations by clustering the MD trajectories of apo-MEIG1, apo-PACRG, and the MEIG1-PACRG complex. This clustering process resulted in 24 distinct MEIG1 conformations (12 from apo-MEIG1 and 12 from the MEIG1-PACRG complex) and 24 PACRG conformations (12 from apo-PACRG and 12 from the MEIG1-PACRG complex). We employed a large number of clusters to mimic the native motions that the proteins would undergo during ligand binding, thereby capturing a comprehensive range of possible conformations. We then performed virtual screening on these 48 protein structures, identifying approximately 60 candidate compounds per structure based on their docking scores, which reflect their binding affinity. In total, 1,395 compounds with high affinity for MEIG1 and 1,377 for PACRG were identified, including some compounds that exhibited binding potential across multiple protein clusters.

By sorting the compounds according to their docking scores, we compiled a list of the top 48 candidate inhibitors for both MEIG1 and PACRG (Tables 1 and 2). Interestingly, none of the top 48 candidate inhibitors for MEIG1 overlapped with those for PACRG, indicating that both proteins bind to very different sets of compounds. Notably, the compounds docked into PACRG exhibited significantly better docking scores compared to those docked into MEIG1, suggesting that the binding sites on PACRG are more favorable for designing a small molecule binder. This highlights PACRG as a potentially more promising target for drug design.

**Table 1.**
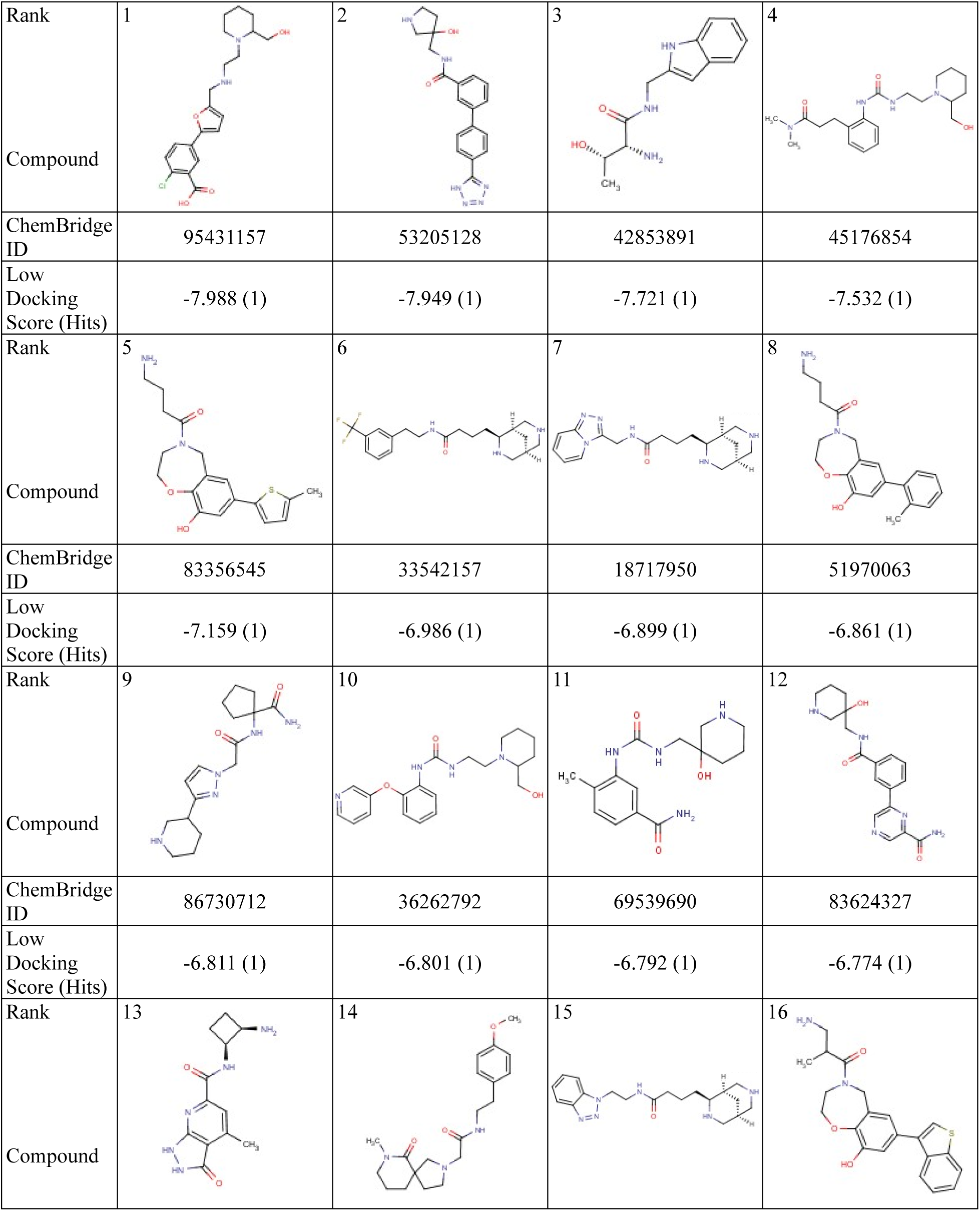
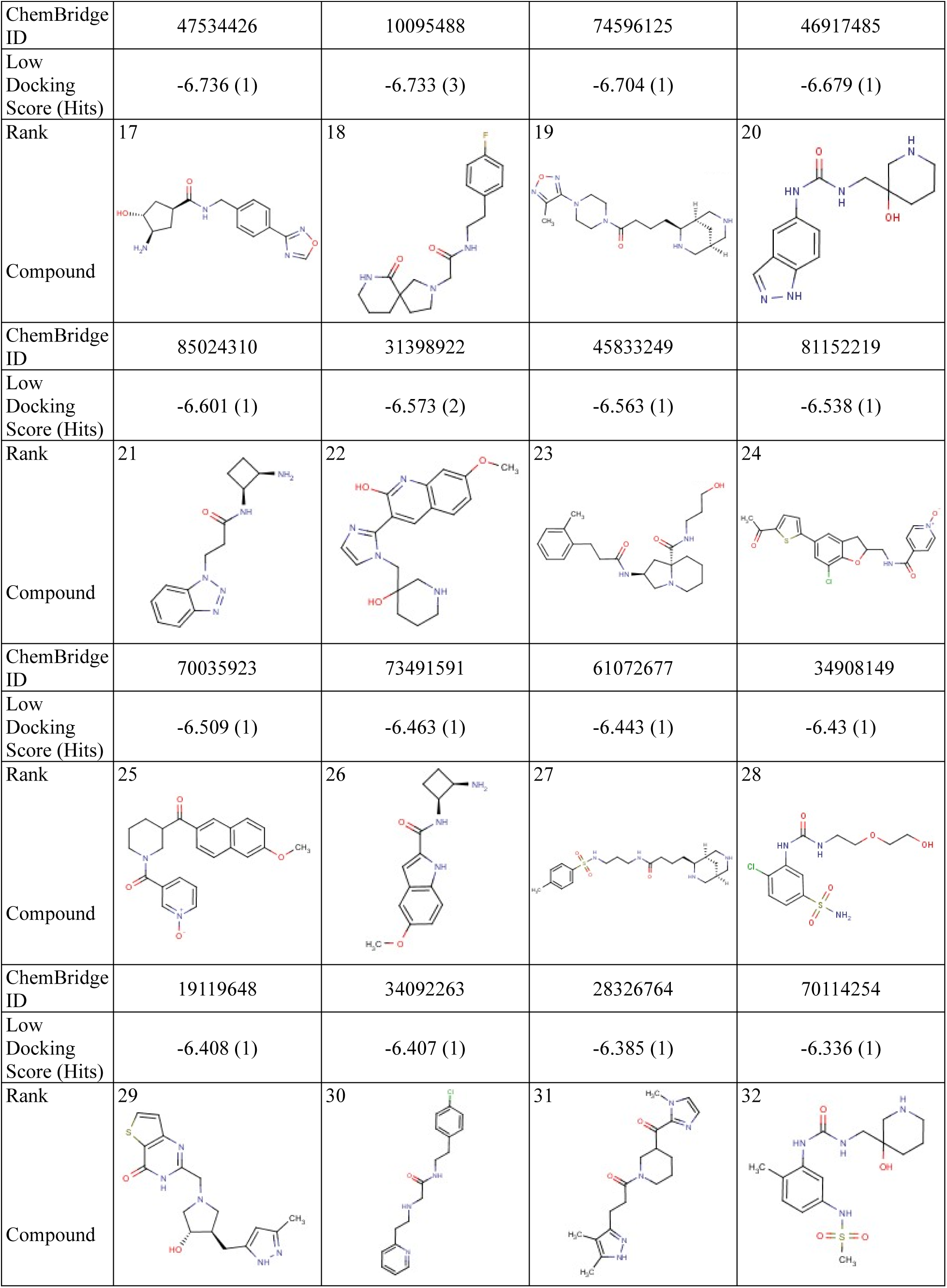
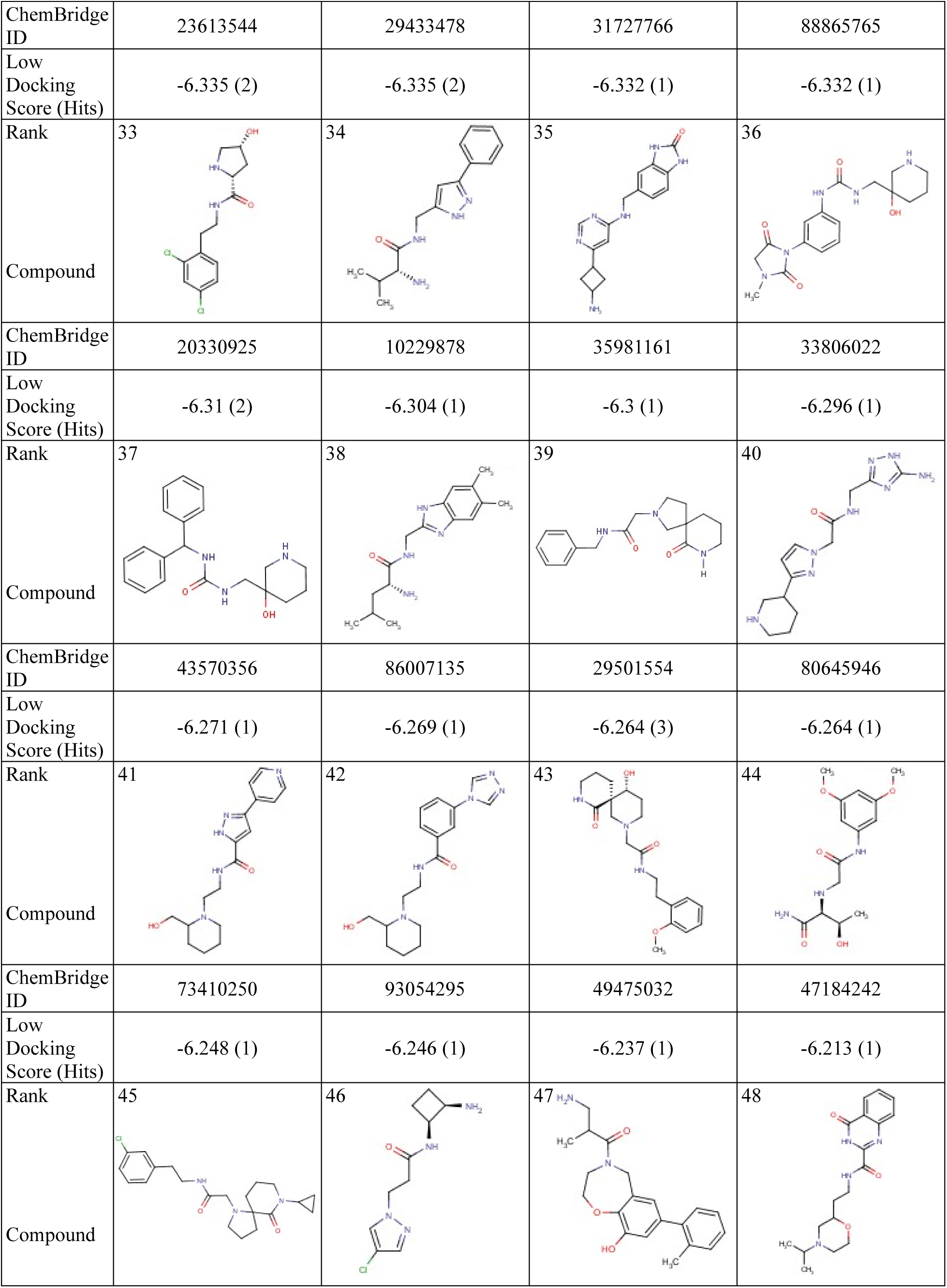
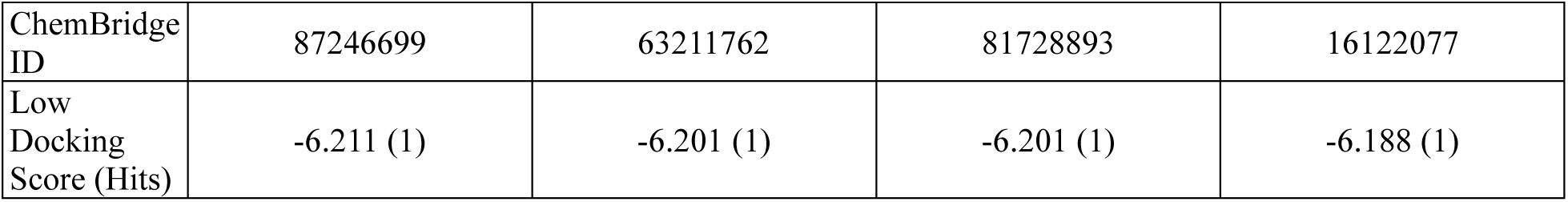
List of top 48 compounds for MEIG1. The top 48 compounds that can bind to MEIG1 were identified through virtual screening and molecular docking. The compounds are ranked by their docking scores. The table displays the chemical structure, Chembridge ID, docking score, and the number of protein clusters where a given compound was identified as a MEIG1 binder (i.e., the number of hits). The ranking of each compound is displayed at the top left of its chemical structure.

**Table 2.**
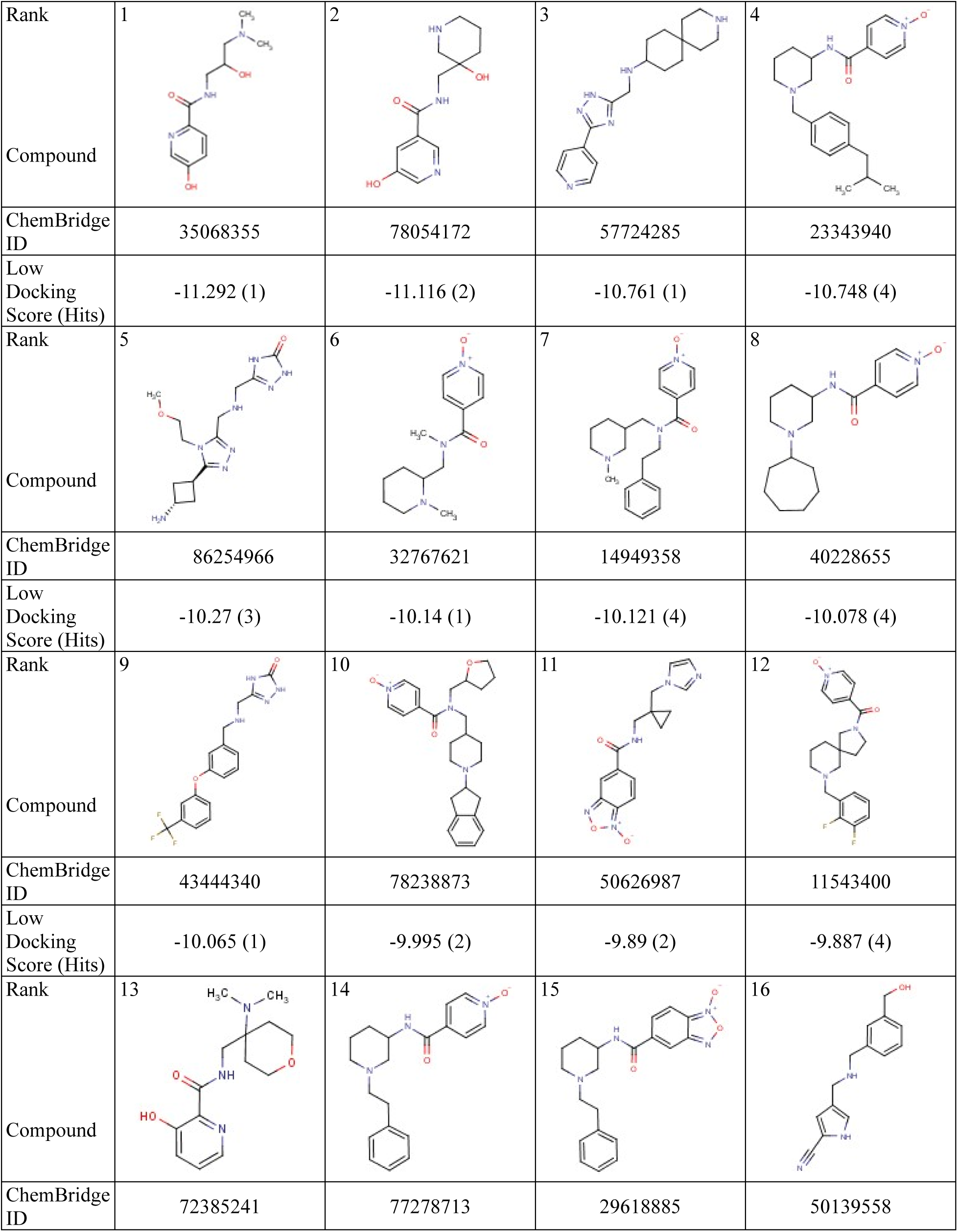
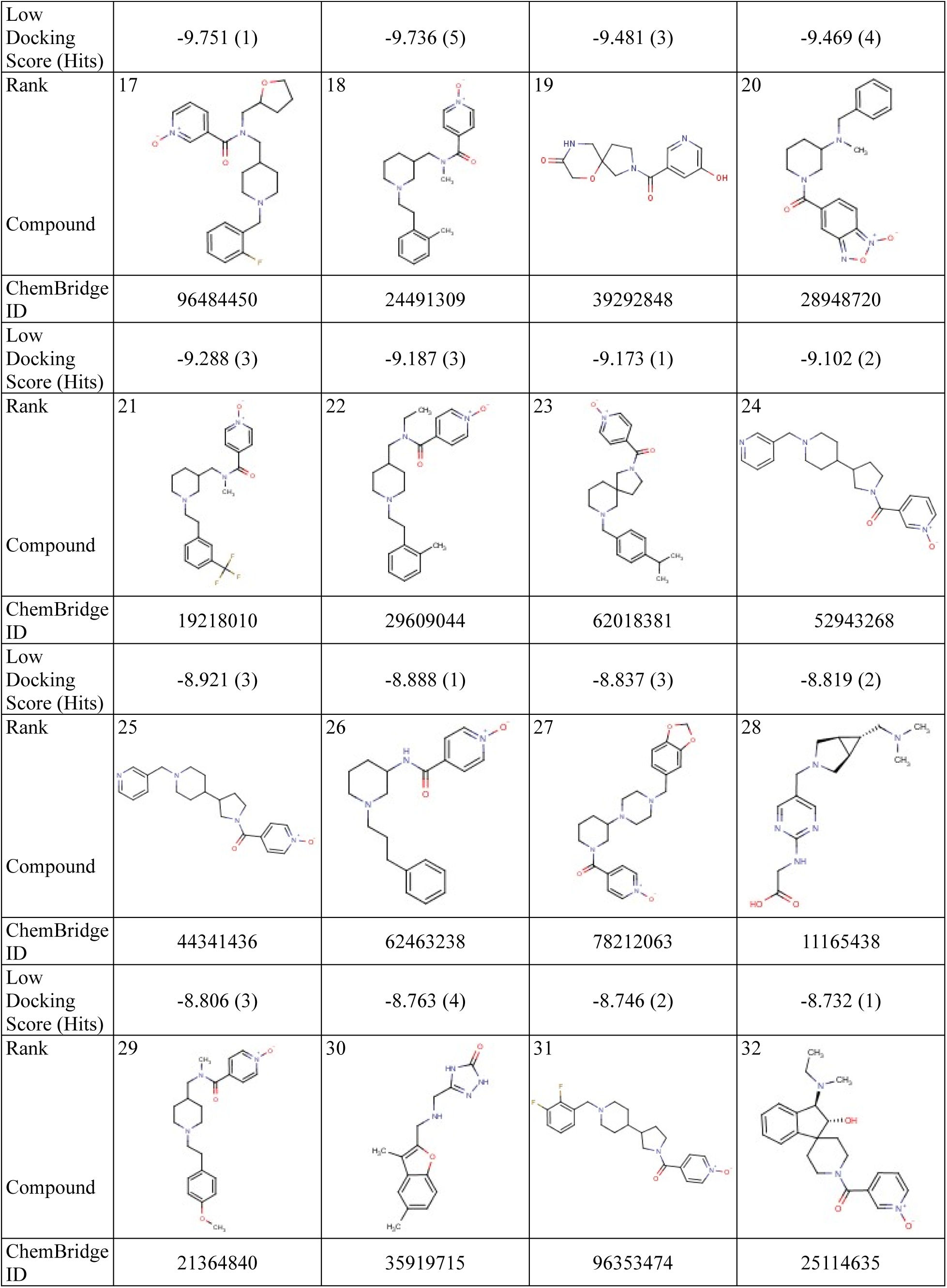
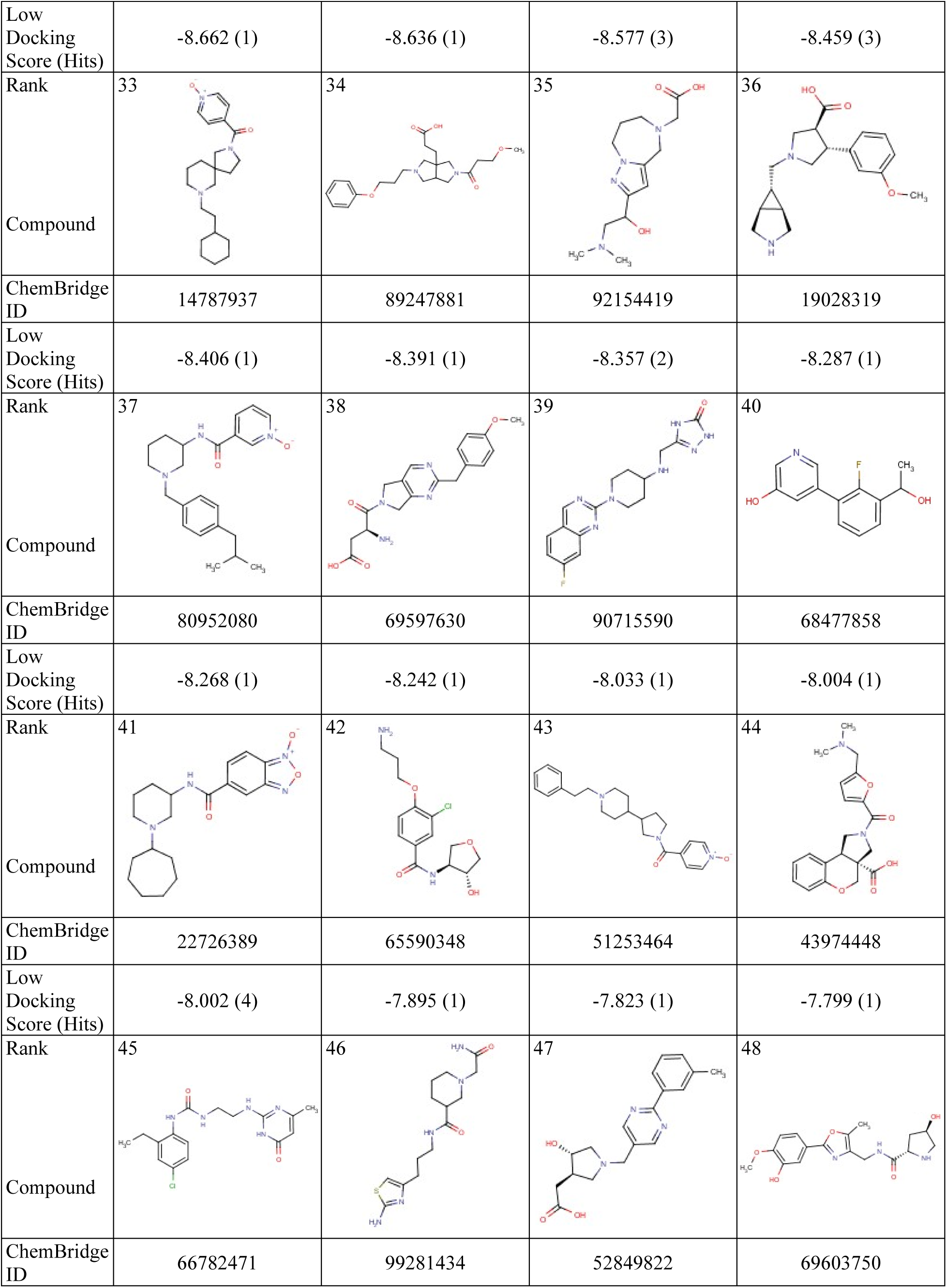
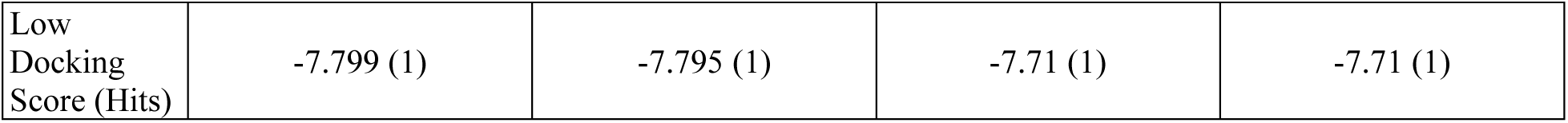
List of top 48 compounds for PACRG. The top 48 compounds that can bind to PACRG were identified through virtual screening and molecular docking. The compounds are ranked by their docking scores. The table displays the chemical structure, Chembridge ID, docking score, and the number of protein clusters where a given compound was identified as a PACRG binder (i.e., the number of hits). The ranking of each compound is displayed at the top left of its chemical structure.

|  |  |  |  |  |
| --- | --- | --- | --- | --- |
| Rank | 1 | 2 | 3 | 4 |
| Compound |  |  |  |  |
| ChemBridge ID | 35068355 | 78054172 | 57724285 | 23343940 |
| Low Docking Score (Hits) | -11.292 (1) | -11.116 (2) | -10.761 (1) | -10.748 (4) |
| Rank | 5 | 6 | 7 | 8 |
| Compound |  |  |  |  |
| ChemBridge ID | 86254966 | 32767621 | 14949358 | 40228655 |
| Low Docking Score (Hits) | -10.27 (3) | -10.14 (1) | -10.121 (4) | -10.078 (4) |
| Rank | 9 | 10 | 11 | 12 |
| Compound |  |  |  |  |
| ChemBridge ID | 43444340 | 78238873 | 50626987 | 11543400 |
| Low Docking Score (Hits) | -10.065 (1) | -9.995 (2) | -9.89 (2) | -9.887 (4) |
| Rank | 13 | 14 | 15 | 16 |
| Compound |  |  |  |  |
| ChemBridge ID | 72385241 | 77278713 | 29618885 | 50139558 |
| Low Docking Score (Hits) | -9.751 (1) | -9.736 (5) | -9.481 (3) | -9.469 (4) |
| Rank | 17 | 18 | 19 | 20 |
| Compound |  |  |  |  |
| ChemBridge ID | 96484450 | 24491309 | 39292848 | 28948720 |
| Low Docking Score (Hits) | -9.288 (3) | -9.187 (3) | -9.173 (1) | -9.102 (2) |
| Rank | 21 | 22 | 23 | 24 |
| Compound |  |  |  |  |
| ChemBridge ID | 19218010 | 29609044 | 62018381 | 52943268 |
| Low Docking Score (Hits) | -8.921 (3) | -8.888 (1) | -8.837 (3) | -8.819 (2) |
| Rank | 25 | 26 | 27 | 28 |
| Compound |  |  |  |  |
| ChemBridge ID | 44341436 | 62463238 | 78212063 | 11165438 |
| Low Docking Score (Hits) | -8.806 (3) | -8.763 (4) | -8.746 (2) | -8.732 (1) |
| Rank | 29 | 30 | 31 | 32 |
| Compound |  |  |  |  |
| ChemBridge ID | 21364840 | 35919715 | 96353474 | 25114635 |
| Low Docking Score (Hits) | -8.662 (1) | -8.636 (1) | -8.577 (3) | -8.459 (3) |
| Rank | 33 | 34 | 35 | 36 |
| Compound |  |  |  |  |
| ChemBridge ID | 14787937 | 89247881 | 92154419 | 19028319 |
| Low Docking Score (Hits) | -8.406 (1) | -8.391 (1) | -8.357 (2) | -8.287 (1) |
| Rank | 37 | 38 | 39 | 40 |
| Compound |  |  |  |  |
| ChemBridge ID | 80952080 | 69597630 | 90715590 | 68477858 |
| Low Docking Score (Hits) | -8.268 (1) | -8.242 (1) | -8.033 (1) | -8.004 (1) |
| Rank | 41 | 42 | 43 | 44 |
| Compound |  |  |  |  |
| ChemBridge ID | 22726389 | 65590348 | 51253464 | 43974448 |
| Low Docking Score (Hits) | -8.002 (4) | -7.895 (1) | -7.823 (1) | -7.799 (1) |
| Rank | 45 | 46 | 47 | 48 |
| Compound |  |  |  |  |
| ChemBridge ID | 66782471 | 99281434 | 52849822 | 69603750 |
| Low<br>Docking<br>Score (Hits) | -7.799 (1) | -7.795 (1) | -7.71 (1) | -7.71 (1) |

Analysis of the top 48 compounds revealed several key features contributing to their high binding affinities: 1) heterocyclic rings (e.g., pyrazole, piperidine, tetrazole, pyridine, and furan), 2) aromatic systems for π-π stacking, and 3) hydrogen bond donors and acceptors (e.g., protonated amine, amide, hydroxyl, and urea groups). A balance of hydrophobic and hydrophilic groups aids both protein-ligand interactions and solubility. Many compounds also include flexible linkers (e.g., alkyl chains), allowing adaptation to dynamic protein conformations. This combination of structural elements makes these compounds strong candidates for targeting the MEIG1-PACRG interaction in drug design.

Our analysis revealed that several compounds exhibited strong affinity for multiple protein clusters (Table 3). Compared to MEIG1, compounds targeting PACRG bound significantly more PACRG clusters. Compounds that bind across multiple MEIG1 conformations generally feature flexible linkers, heterocyclic scaffolds, and polar functional groups, enabling adaptation to different binding environments. However, the limited number of compounds that can bind to multiple MEIG1 clusters suggests that MEIG1’s binding pockets are more structurally restrictive. In contrast, PACRG’s binding sites appeared more accommodating to attract a variety of compounds. Most compounds that bind to multiple PACRG clusters contain aromatic rings and polar groups that facilitate electrostatic and π-π stacking interactions across diverse PACRG conformations. This flexibility of PACRG binding pockets indicates that it may be a more versatile drug target, with its broad compound compatibility underscoring potential for disrupting the MEIG1-PACRG interaction.

**Table 3.**
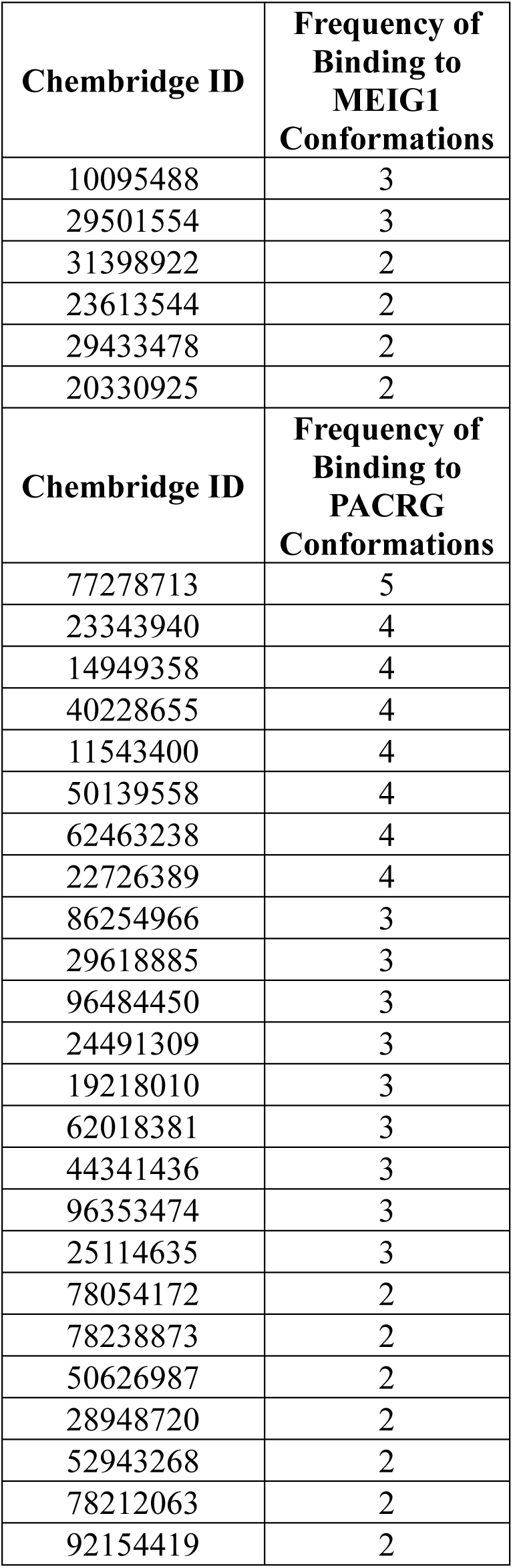
Lists of binding frequency of compounds with potential affinity for MEIG1 and PACRG. Compounds that can bind to more than one protein cluster are shown. The ChemBridge ID for each compound is listed along with the number of different protein clusters it bound to.

| **Chembridge ID** | **Frequency of Binding to MEIG1 Conformations** |
| --- | --- |
| 10095488 | 3 |
| 29501554 | 3 |
| 31398922 | 2 |
| 23613544 | 2 |
| 29433478 | 2 |
| 20330925 | 2 |
| **Chembridge ID** | **Frequency of Binding to PACRG Conformations** |
| 77278713 | 5 |
| 23343940 | 4 |
| 14949358 | 4 |
| 40228655 | 4 |
| 11543400 | 4 |
| 50139558 | 4 |
| 62463238 | 4 |
| 22726389 | 4 |
| 86254966 | 3 |
| 29618885 | 3 |
| 96484450 | 3 |
| 24491309 | 3 |
| 19218010 | 3 |
| 62018381 | 3 |
| 44341436 | 3 |
| 96353474 | 3 |
| 25114635 | 3 |
| 78054172 | 2 |
| 78238873 | 2 |
| 50626987 | 2 |
| 28948720 | 2 |
| 52943268 | 2 |
| 78212063 | 2 |
| 92154419 | 2 |

### Interactions between MEIG1 and potential binders

We screened compounds that bind along the MEIG1-PACRG interaction interface, focusing on areas of MEIG1 that interact with PACRG in their complex. The top 48 MEIG1 compounds exhibited strong interactions with key residues near this interface, including Y25, Q29, W50, K57, Y67, Y68, Y69, N70, K71, and E74 (Figure 2). These residues are crucial for mediating the interaction between MEIG1 and PACRG.

**Figure 2.**
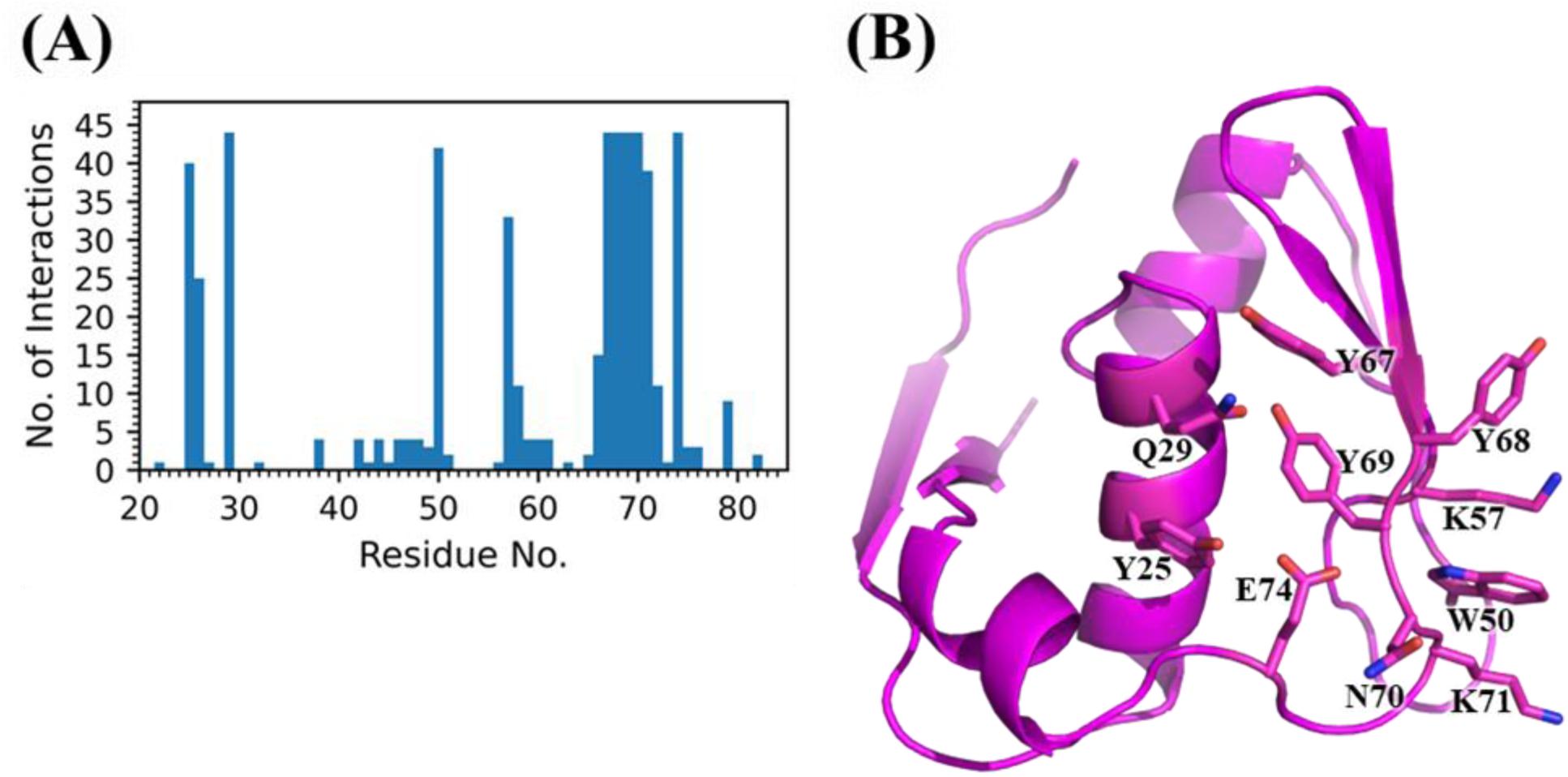
Interaction analysis for MEIG1-ligand complexes. (A) The interaction frequency of MEIG1 residues with the top 48 compounds is shown. The x-axis represents the MEIG1 residue number, and the y-axis shows the number of compounds interacting within a 5 Å from each residue. (B) The MEIG1 structure displays key residues (Y25, Q29, W50, K57, Y67, Y68, Y69, N70, K71, and E74) that frequently interact with compounds.

Compounds binding to MEIG1 generally followed a common binding mode: 1) hydrogen bond donors in the compounds interacted with N70 or E74, 2) central aromatic groups formed π-π stacking interactions with W50 or Y68, and occasionally with Y67 and Y69, and 3) an amide NH group centrally positioned in the compounds typically formed a hydrogen bond with the backbone oxygen of Y68 (Figure 3A). This interaction allowed the surrounding atoms in the compound to bind residues 67-70, creating a central interaction hub that provided multiple contact points and enhanced protein-ligand stability.

**Figure 3.**
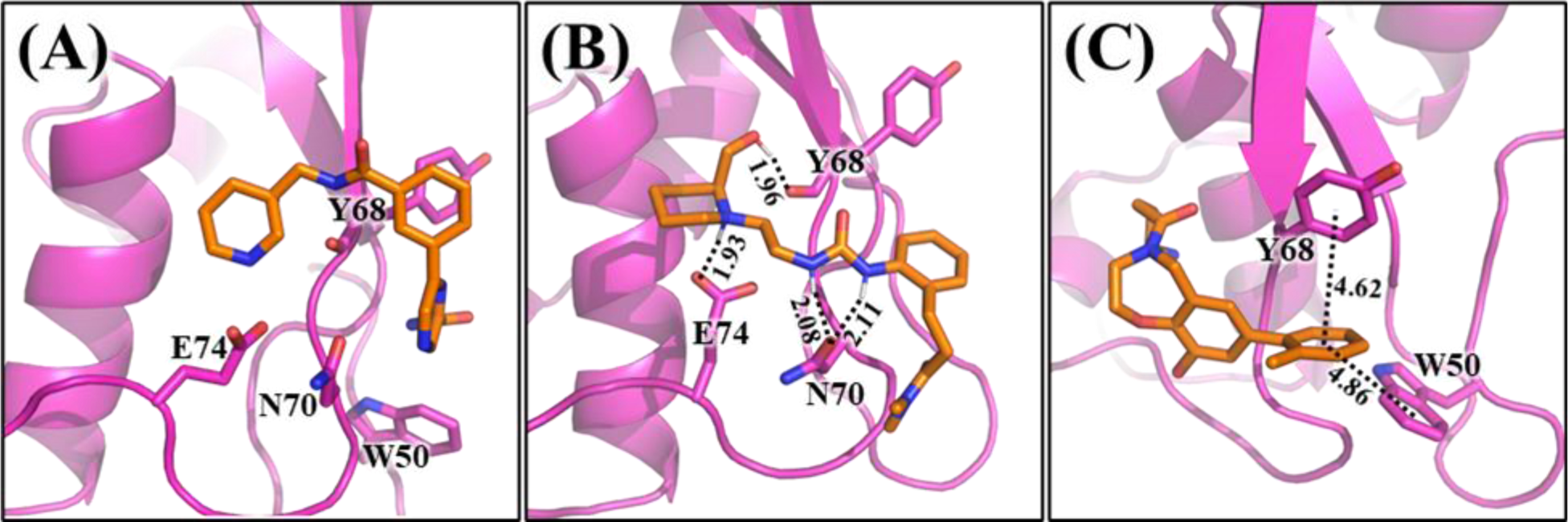
Representative compound binding modes of MEIG1. (A) Binding example of compound (ChemBridge ID: 83624327) to MEIG1, showing key interactions such as hydrogen bonds and aromatic stacking that illustrate the most common binding mode observed in MEIG1-ligand complexes. (B) Binding example of compound (ChemBridge ID: 45176854) to MEIG1, with hydrogen bonds formed between MEIG1 residues Y68 (backbone carbonyl oxygen), E74 (side chain carboxylate oxygen), and N70 (side chain carbonyl oxygen) and the compound. (C) Binding example of compound (ChemBridge ID: 51970063) to MEIG1, showing aromatic interactions between W50 and Y68 side chains and aromatic groups on the compound. Distances in angstroms (Å) between the centers of W50 and Y68 aromatic rings and those on the compound highlight π-π stacking interactions.

Hydrogen bonding is crucial in protein-compound interactions, with key residues Y68, N70, and E74 consistently forming hydrogen bonds with compounds (Figure 3B). These residues acted as hydrogen bond acceptors, typically interacting with nitrogen-containing groups from compounds, including amines, amides, and heterocyclic structures. E74 formed most hydrogen bonds, particularly with protonated piperidine groups or primary amines on alkyl chains. Both N70 and the backbone carbonyl oxygen of Y68 bonded with neutral amines, amides, or hydroxyl groups. Occasionally, Y25, Q29, K57, K71, and the backbone carbonyl oxygen Y69 also formed hydrogen bonds, but these were secondary to those involving Y68, N70, and E74.

Aromatic interactions were also prevalent, particularly between the sidechains of W50 and Y68 and aromatic substituents on the compounds, such as benzene rings or nitrogen-containing heterocycles (Figure 3C). These π-π stacking interactions occurred in configurations such as offset face-to-face (parallel displaced), edge-to-face (T-shaped), and face-to-face (parallel stacking), with distances typically ranging from 4.0 to 6.0 Å. Additionally, van der Waals and hydrophobic interactions, involving residues like W50, Y67, Y68, Y69, and alkyl side chains of K57 and K71, further stabilized the binding.

### Interactions between PACRG and potential binders

We screened compounds that could bind along the PACRG-MEIG1 interaction interface, targeting regions of PACRG involved in the complex with MEIG1. The top 48 PACRG compounds exhibited strong interactions with residues K93, W96, E99, I100, E101, D133, and H137, all located near the PACRG-MEIG1 interaction interface (Figure 4).

**Figure 4.**
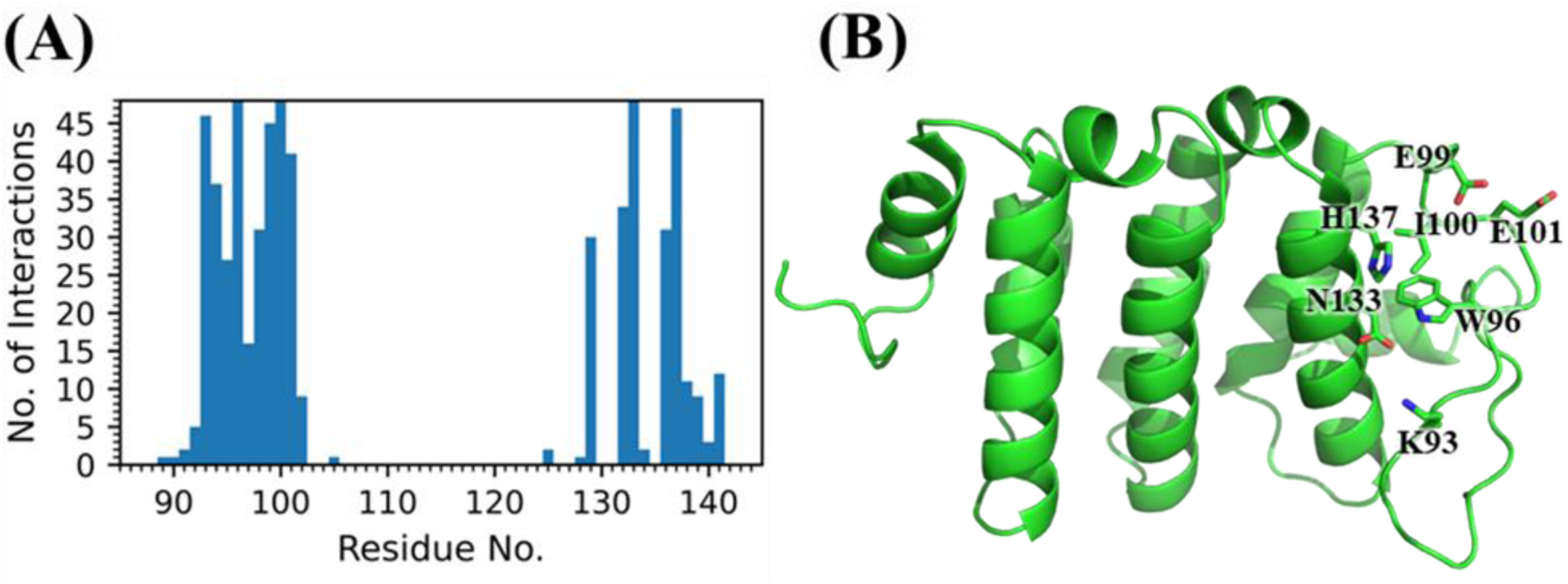
Interaction analysis for PACRG-ligand complexes. (A) The interaction frequency of PACRG residues with the top 48 compounds is shown. The x-axis represents the PACRG residue number, and the y-axis shows the number of compounds interacting within a 5 Å from each residue. (B) The PACRG structure displays key residues (K93, W96, E99, I100, E101, D133, and H137 that frequently interact with compounds.

In most PACRG-compound complexes, compounds form hydrogen bonds with K93 and E101 at opposite ends. The central region of the compound, often featuring aromatic rings or nitrogen-containing heterocycles, interacts with W96, I100, and H137 (Figure 5A). E99 and N133 also participate in hydrogen bonding, though these interactions occur less frequently than those with K93 and E101.

**Figure 5.**
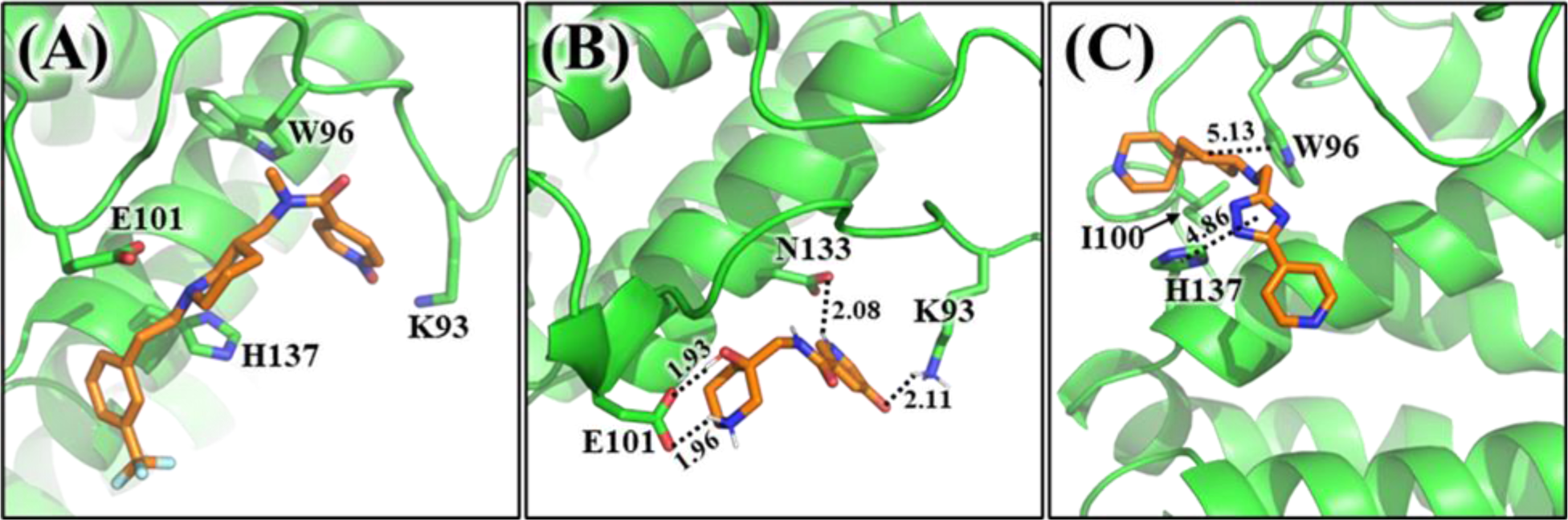
Representative compound binding modes of PACRG. (A) Binding example of compound (ChemBridge ID: 19218010) to PACRG, showing key interactions such as hydrogen bonds and hydrophobic interactions that illustrate the most common binding mode observed in PACRG-ligand complexes. (B) Binding example of compound (ChemBridge ID: 78054172) to PACRG, with hydrogen bonds formed between PACRG residues K93 (side chain protonated amine), E101 (side chain carboxylate oxygen), and the compound. (C) Binding example of compound (ChemBridge ID: 57724285) to PACRG, showing aromatic and van der Waals interactions between W96 and H137 side chains and aromatic and hydrophobic groups on the compound. Distances in angstroms (Å) between the centers of W96 and H137 aromatic rings and those on the compound highlight π-π stacking interactions.

Hydrogen bonding dominated PACRG-compound interactions, particularly through K93 and E101 (Figure 5B). The side chain of K93, with its protonated amine group, acted as a hydrogen bond donor, typically forming interactions with oxygen-containing groups on compounds such as carboxylic acids, hydroxyl groups, triazole-3-one, benzofuroxan, and most commonly, isonicotinamide 1-oxide groups. E101, as a hydrogen bond acceptor, interacted with protonated nitrogen groups on the compounds, especially piperidine, as well as amines. These interactions were critical for anchoring compounds within the PACRG binding pocket, suggesting that targeting K93 and E101 is a key strategy for disrupting the PACRG-MEIG1 interaction.

Aromatic interactions were observed between W96 and H137 with the aromatic rings of compounds, such as benzene or heterocyclic groups (Figure 5C). Compounds positioned between W96 and H137 formed edge-to-face or parallel stacking interactions with both residues, further stabilizing the binding. In addition, I100 also contributed to van der Waals and nonpolar interactions, interacting with both aromatic and alkyl chains on the compounds, thereby reinforcing the overall binding stability.

The binding pocket of PACRG is more concave and well-defined compared to that of MEIG1, providing a favorable fit for compounds. Key residues, such as K93 and E101, exhibit flexibility within this pocket, enhancing its capacity to accommodate diverse compound structures and improve docking scores. PACRG’s ability to bind various compounds across multiple conformations highlights its potential as a promising target for drug design, as its adaptable binding pocket can interact effectively with a broader range of inhibitors.

### Molecular dynamics simulations of top MEIG1 binders

To assess the dynamics of the MEIG1-binding compounds, we conducted 100 ns conventional MD simulations on the 10 complexes with the best docking scores. The simulations indicated that most compounds remained tightly bound within the MEIG1 binding pocket, preserving all critical interactions for the entire duration of the simulations. One compound exhibited partial unbinding, while some compounds fluctuated within the pocket, repositioning their interactions (Figure 6). Hydrogen bonds between E74 and compounds (Figure 3B) remained consistently stable throughout the simulations, occasionally reorienting to interact with alternative substituents on the compound. In contrast, hydrogen bonds involving the N70 sidechain displayed greater variability; they frequently dissociated but sometimes reformed with different groups on the compound. Interestingly, N70 occasionally shifted roles, acting as a hydrogen bond donor via its sidechain nitrogen instead of its more typical role as a hydrogen bond acceptor through the sidechain oxygen. The backbone oxygen of Y68 exhibited a high tendency to dissociate, with minimal reestablishment of interactions during the simulations. Our results underscore the pivotal role of hydrogen bonding with E74 as the primary interaction anchoring compounds within the MEIG1 binding pocket.

**Figure 6.**
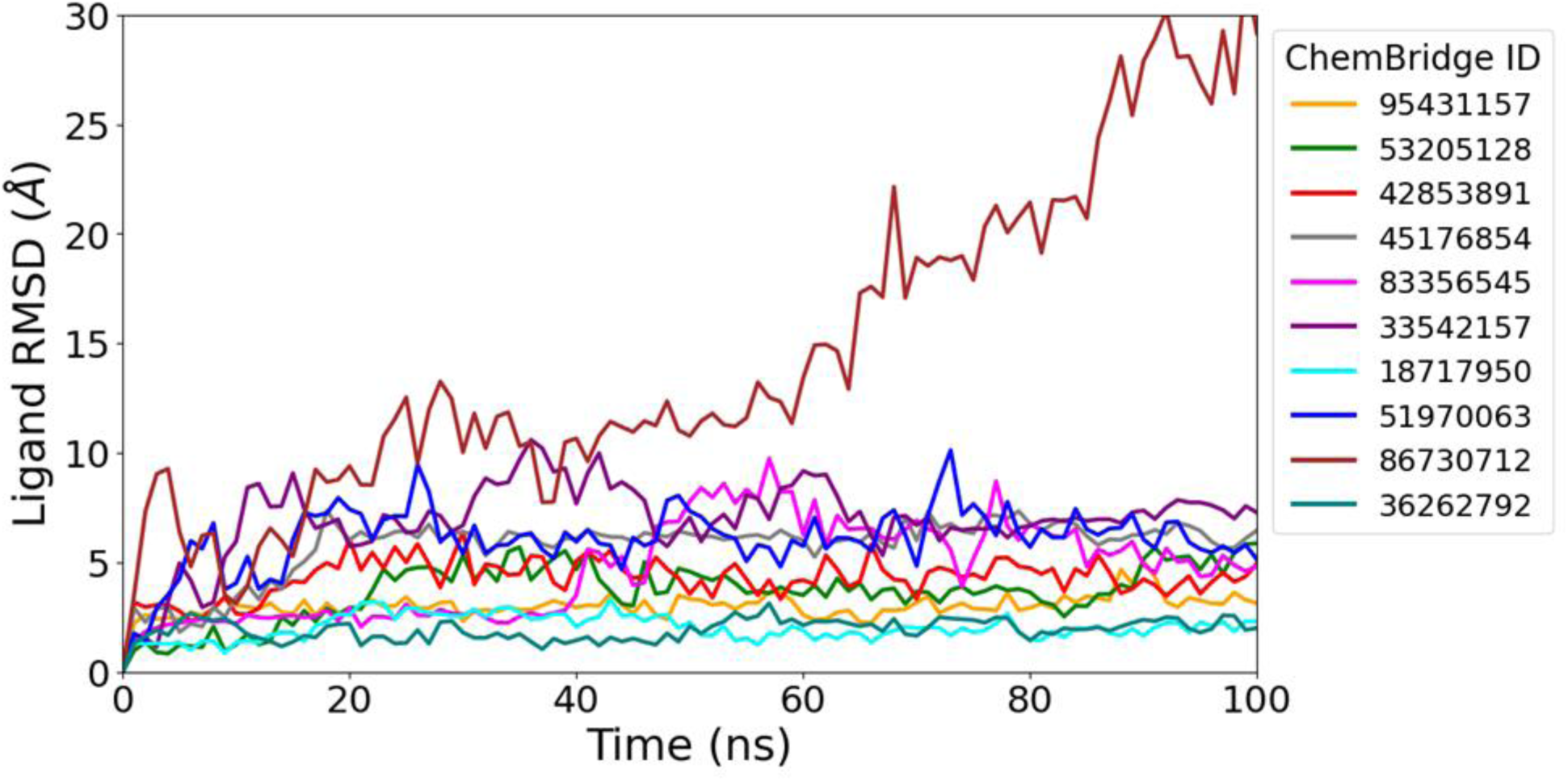
RMSD for the top 10 MEIG1-binding compounds during 100 ns MD simulations. The RMSD was calculated using the compound heavy atoms relative to the initial docked conformation. Each trajectory is labeled with the corresponding ChemBridge ID.

In addition to hydrogen bonding, π-π stacking interactions were identified as critical stabilizing forces for compound binding. Residues W50 and Y68 (Figure 3C) consistently formed strong and persistent interactions with compounds during the simulations. These interactions often intensified as the aromatic, amide, and heterocyclic groups on the compounds aligned closely with the sidechains of W50 and Y68. Additional contributions from Y67 and Y69 provided supplementary π-π stacking interactions. The ability of the compounds to reorient within the binding pocket enabled them to optimize their stacking arrangements, thereby significantly enhancing the stability of the protein-ligand complexes. These findings emphasize the dominant role of π-π stacking, particularly with W50 and Y68, in maintaining compound binding within the MEIG1 pocket, compensating for the variability or dissociation observed in some hydrogen bonds, such as those involving Y68 and N70.

ChemBridge ID 33542157 provides a detailed example of compound reorientation involving hydrogen bonding and π-π stacking during simulations (Figure 7). In the docking conformation, the diazabicyclo scaffold from the compound established hydrogen bonds with the sidechain oxygen of E74 and the backbone oxygen of Y68. Its amide group interacted with N70, while an edge-to-face π-π stacking interaction between the trifluoromethylphenyl group and W50 occurred (Figure 7A). As the simulation progressed, the compound underwent significant reorientation within the MEIG1 binding pocket. The trifluoromethylphenyl group shifted, ultimately forming face-to-face stacking with Y68 and maintaining interactions with W50 (Figure 7B). The hydrogen bonds involving Y68 and N70 dissociated, while E74 consistently maintained its interactions, reinforcing its critical role in complex stability (Figure 7C). Measurements of centroid distances and interplanar angles (Figure 7D and 7E) demonstrated how the aromatic trimethylphenyl ring of the compound dynamically fluctuated between edge-to-face stacking (∼6 Å, 60°–90°) and face-to-face stacking (∼4 Å, 0°–30°) with Y68 and W50. These changes indicated a transition in the binding mode, evolving from initial hydrogen bond dominance to increasing stabilization through π-π stacking interactions. Hydrogen bonding with E74 and π-π stacking involving W50 and Y68 was pivotal in maintaining the structural integrity of the MEIG1-compound complex throughout the simulation.

**Figure 7.**
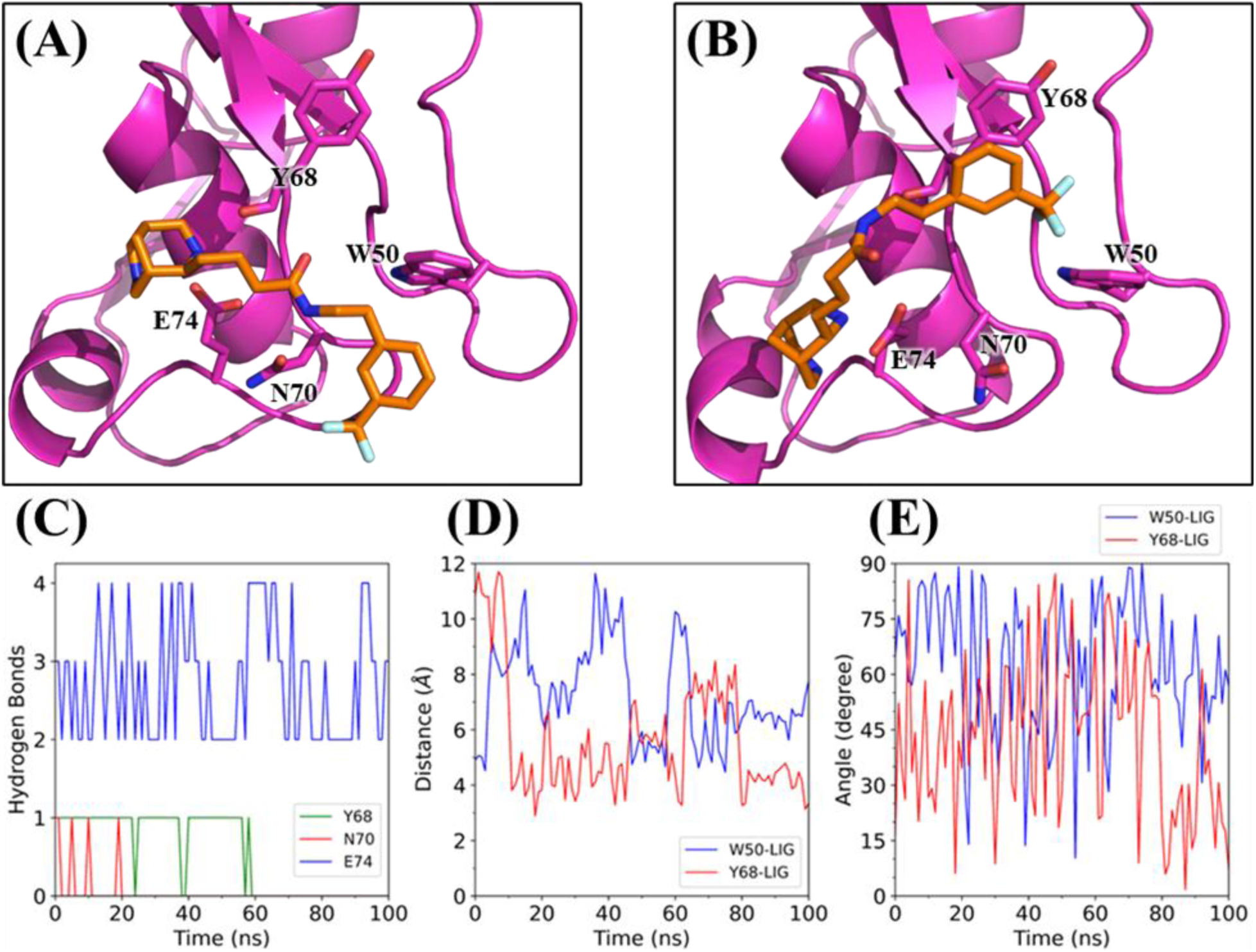
Representative interaction analysis for MEIG1-compound complexes from MD simulations. (A) The initial docked conformation of the compound (ChemBridge ID: 33542157) bound to MEIG1 highlights key hydrogen bonds with Y68 (backbone carbonyl oxygen), N70 (side chain carbonyl oxygen), and E74 (side chain carboxylate oxygen), as well as π-π stacking interactions with W50. (B) The compound conformation extracted after the 100-ns MD simulation reveals a transition to face-to-face π-π stacking with Y68 and edge-to-face stacking with W50. (C) Time evolution of hydrogen bonds between MEIG1 residues Y68 (green), N70 (red), and E74 (blue) and the compound. (D) Distances between the aromatic centroids of MEIG1 residues W50 (blue) and Y68 (red) and the compound’s aromatic trifluoromethylphenyl group throughout the simulation. (E) Angles between the planes of MEIG1 residues W50 (blue) and Y68 (red) and the compound’s aromatic group throughout the simulation.

### Molecular dynamics simulations of top PACRG binders

We performed 100 ns conventional MD simulations on the top 10 PACRG-binding compounds to investigate their dynamics. All compounds remained stably bound within the PACRG pocket, maintaining low RMSD values throughout the simulations (Figure 8). However, hydrogen bond interactions showed significant variability, highlighting the dynamic nature of the residues involved. The most consistent interaction was with E101 (Figure 5B), which remained stable throughout the simulations for several compounds. In many cases, dissociation of E101 was followed by compensatory hydrogen bonds involving E99. On the other hand, K93, located on the opposite side of the pocket, displayed intermittent bonding, dissociating early in the simulations and rarely reforming due to its reorientation away from the binding pocket. N133, positioned at the base of the pocket’s concavity, exhibited stable hydrogen bonding in some trajectories but was absent in others. These variations suggest that PACRG residues are more flexible compared to the rigid interactions observed with MEIG1. Despite these fluctuations, the adaptability of the hydrogen bonding network, especially the stability of the E101 interaction, was crucial in maintaining the overall stability of the PACRG-compound complexes.

**Figure 8.**
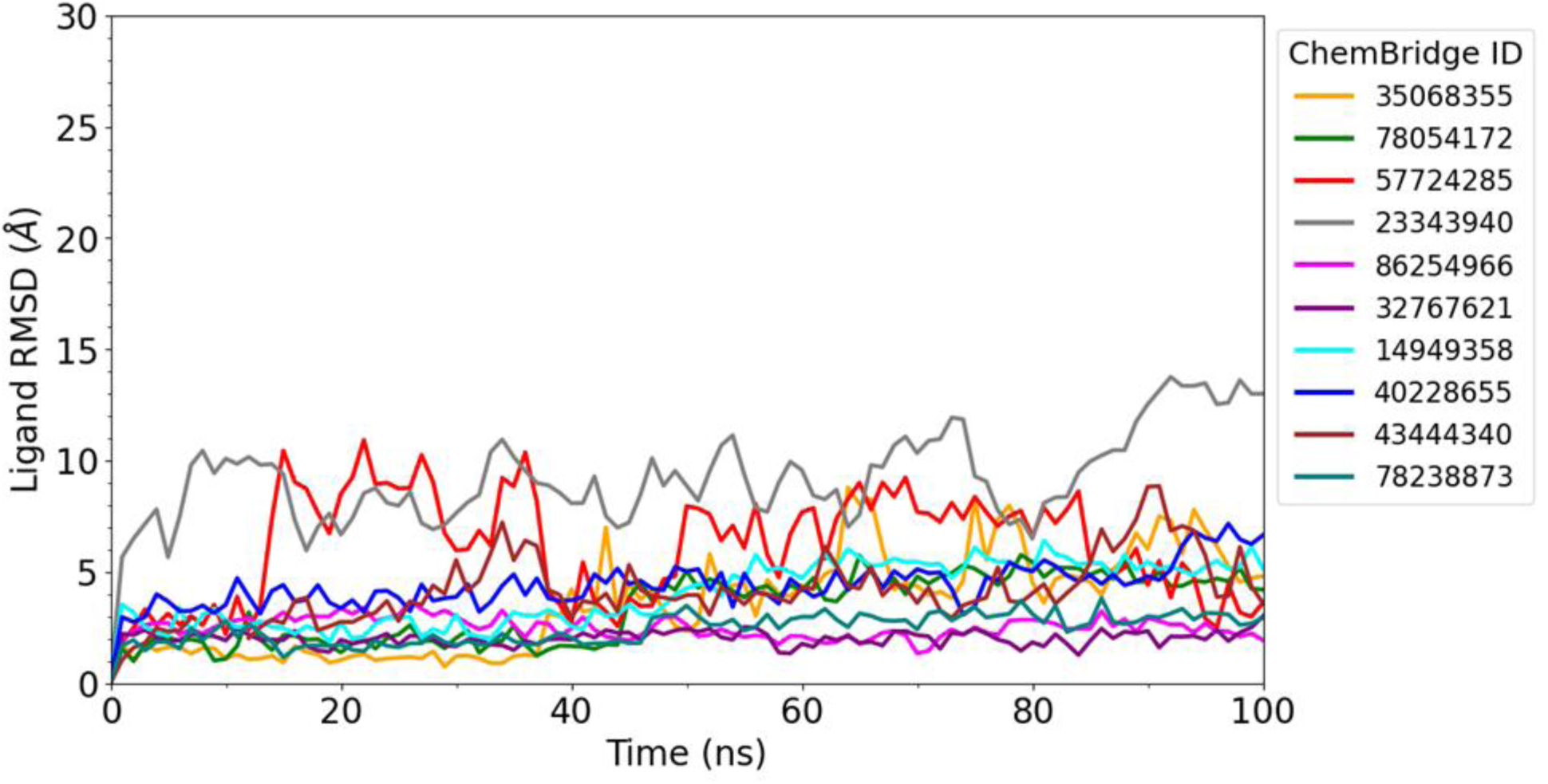
RMSD for the top 10 PACRG-binding compounds during 100 ns MD simulations. The RMSD was calculating using the compound heavy atoms relative to the initial docked conformation. Each trajectory is labeled with the corresponding ChemBridge ID.

π-π stacking interactions with residues W96 and H137 (Figure 5C) played a central role in stabilizing PACRG-compound complexes. The interactions frequently transitioned between face-to-face and edge-to-face configurations with W96 and H137, preserving centroid distances and interplanar angles favorable for strong π-π stacking. These interactions compensated for the variability of hydrogen bonds and provided additional stabilization of the compounds within the binding pocket. Notably, W96 and H137 effectively bound the compounds, particularly while the hydrogen bonds of K93 or N133 dissociated. I100, located near W96 and H137, contributed to additional compound binding through hydrophobic interactions with the same compound aromatic groups that interacted with W96 and H137. These findings emphasize that the stability of PACRG-compound complexes depends significantly on the cooperative effects of aromatic interactions. These interactions provided a robust stabilizing framework, compensating for the fluctuating nature of individual hydrogen bonds and underscoring their importance in maintaining complex integrity.

ChemBridge ID 78238873 provides an example to illustrate the dynamic interactions in protein-compound complexes (Figure 9). The initial docked conformation (Figure 9A) revealed hydrogen bonds between the compound and residues K93 and E101, alongside π-π stacking interactions between the compound’s isonicotinamide 1-oxide group and the aromatic sidechains of W96 and H137. By the end of the simulation (Figure 9B), the hydrogen bond with E101 remained stable, while the hydrogen bond with K93 dissociated early as K93 reoriented away from the compound. E101 maintained a consistent hydrogen bond throughout the simulation, as shown in Figure 9C. Meanwhile, the π-π stacking interactions involving W96 and H137 fluctuated dynamically, with the compound’s aromatic group reorienting to maintain close associations with both residues. Centroid-centroid distances and interplanar angles (Figure 9D and 9E) revealed fluctuating π-π stacking configurations with W96 and H137. Additionally, H137 was stabilized by nonpolar interactions, as the indane, piperidine, and tetrahydro-2-furanyl groups of the compound formed van der Waals contacts with the surrounding hydrophobic environment. These results emphasize the role of aromatic and van der Waals interactions, particularly with W96 and H137, in offsetting the loss of less stable hydrogen bonds, such as those involving K93.

**Figure 9.**
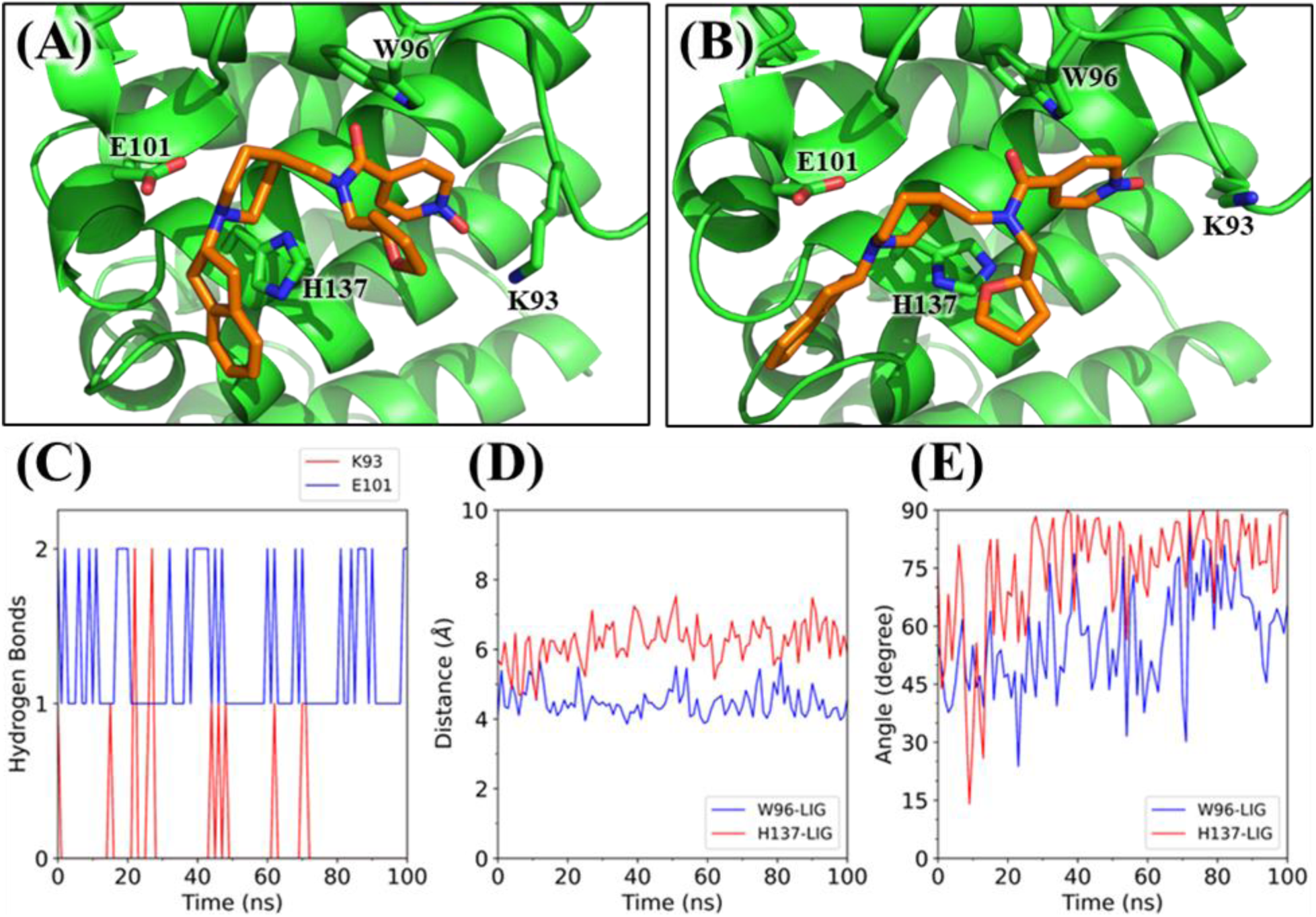
Representative interaction analysis for PACRG-compound complexes from MD simulations. (A) The initial docked conformation of the compound (ChemBridge ID: 78238873) bound to PACRG highlights key hydrogen bonds with K93 (sidechain amino group) and E101 (sidechain carboxylate oxygen), as well as π-π stacking interactions with W96 and H137. (B) The compound conformation extracted from the 100-ns MD simulation reveals the loss of the hydrogen bond with K93 while retaining the hydrogen bond with E101 and the aromatic interactions with W96 and H137. (C) Time evolution of hydrogen bonds between PACRG residues K93 (red) and E101 (blue) and the compound. (D) Distances between the aromatic centroids of PACRG residues W96 (blue) and H137 (red) and the compound’s aromatic isonicotinamide 1-oxide group throughout the simulation. (E) Angles between the planes of PACRG residues W96 (blue) and H137 (red) and the compound’s aromatic group throughout the simulation.

## Conclusions

In this study, we employed virtual screening, molecular docking, and MD simulations to identify small molecules capable of binding to MEIG1 or PACRG, with the goal of disrupting the MEIG1-PACRG interaction — a key regulator of spermiogenesis and male fertility. Virtual screening and molecular docking were performed using diverse protein conformations generated from MD simulations, incorporating protein flexibility into the screening process. Our comprehensive computational analyses identified top candidate compounds with strong binding affinities to MEIG1 and PACRG, specifically targeting the interface where these two proteins interact. Virtual screening revealed distinct sets of compounds for MEIG1 and PACRG, with PACRG emerging as a more favorable drug target due to its better docking scores and more flexible binding pockets. Key residues involved in compound binding were identified, including W50, Y68, N70, and E74 of MEIG1, and K93, W96, E101, and H137 of PACRG. MD simulations elucidated the dynamics of the protein-compound complexes and the significance of specific molecular interactions in maintaining binding affinity. For MEIG1, hydrogen bonding with E74 and π-π stacking interactions with W50 and Y68 were pivotal in maintaining compound stability. For PACRG, stabilization of the protein-compound complex was primarily driven by consistent hydrogen bonding with E101 and π-π stacking interactions with W96 and H137, which were complemented by hydrophobic contacts with I100. Our work not only identified potential candidate compounds for further experimental validation but also offered valuable insights into the molecular mechanisms of compound binding that could disrupt MEIG1-PACRG interactions. These findings are anticipated to contribute significantly to the development of innovative non-hormonal male contraceptive methods, providing a new avenue for addressing male reproductive health.

## Acknowledgments

We express our gratitude to the Wayne State University High-Performance Computing Center for supporting our virtual screening, molecular docking and MD simulations. This research is supported by the Wayne State University Start-up fund to Y.M.H, and the National Institutes of Health under award numbers RO1-HD105944, RO1 HD114311, and R21 HD107579 to Z.Z.

## Data availability statement

The results of virtual screening and MD simulations are freely available at https://doi.org/10.5281/zenodo.14284094.

